# Introns are derived from transposons

**DOI:** 10.1101/2023.02.21.529479

**Authors:** Scott O. Rogers, Arnold J. Bendich

## Abstract

Introns and transposons exhibit many similar features, but the connections between them have yet to be firmly established. Group I introns have commonalities with DNA transposons, while group II introns share many features with retrotransposons. Here, we report the results of an analysis of 214 introns (including group I, group II, group III, twintrons, spliceosomal, and archaeal introns) from members of seven major taxa (within Eukarya, Bacteria, and Archaea) that all have direct repeats at or near both exon/intron borders, indicating that they were inserted via transposition events. Border sequence analysis indicates that after splicing, most mature transcripts would be functionally compromised because they do not restore the DNA sequence information before intron insertion. Transposons and introns thus appear to be members of a diverse assemblage of parasitic mobile genetic elements that secondarily may benefit their host cell and have expanded greatly in eukaryotes from their presumed prokaryotic ancestors.

**Author Summary:** Introns are found in all domains of life. While they are limited in prokaryotes, they have greatly expanded in number and diversity in eukaryotes. We found direct repeat sequences at or near both exon/intron borders for all 214 introns analyzed among eukaryotes, bacteria, and archaea. We infer that all introns were inserted into genes via transposon-like mechanisms and are members of a large family of mobile genetic elements.

## Introduction

Cellular genes contain DNA sequences that encode RNA and protein products. In order to benefit the cell, those sequences typically must be contiguous and uninterrupted by extraneous DNA sequences that would compromise function. Introns do, however, separate the functional parts of genes and are typically removed from the RNA transcripts allowing the RNA to contribute to cellular activities. Introns are present in all domains of life and are identified by comparing the intron-containing gene with the same gene present in an organism lacking the intron [1–4]. The fact that one strain of a species can carry an intron that is absent from another strain indicates that introns were mobile in the past and may still be mobilized, a property shared with mobile genetic elements (MGEs) that will be considered below. Although the origin of introns is enigmatic [5–8], they have been classified according to the way in which they are removed from gene transcripts (Fig 1).

**Figure 1.**
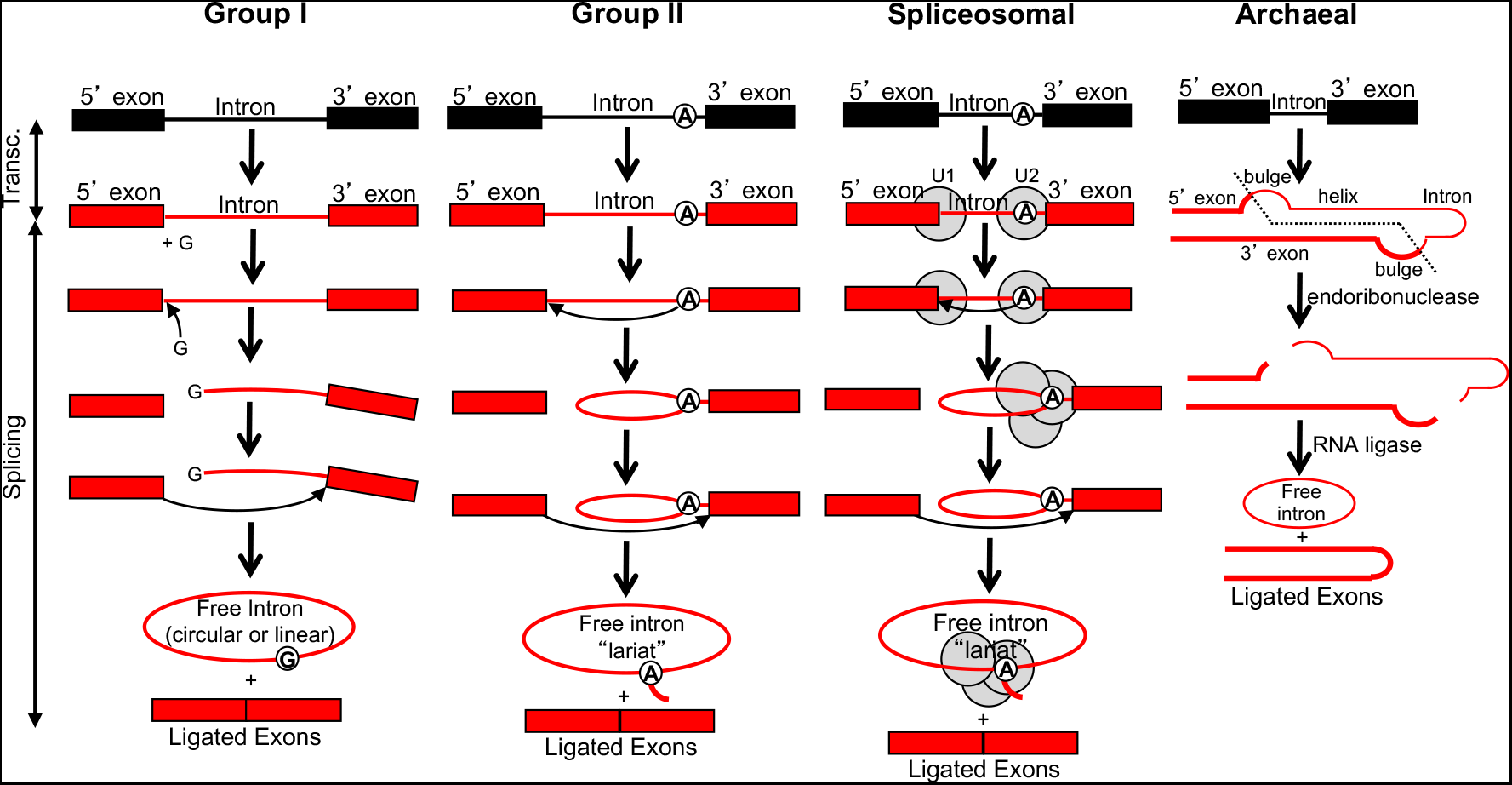
Integration and splicing of intron types. Group I introns are self-splicing: they are ribozymes containing sequences that can cause cleavage and ligation reactions. Two transesterification reactions occur. The first is a nucleophilic attack on the 5’ splice border by the 3’ hydroxyl of the free guanosine (G) held in pairing region P7, which separates the 5’ exon from the intron, leaving the G covalently bonded to the 5’ end of the intron. The second nucleophilic attack is initiated by the 3’ hydroxyl at the 3’ end of the 5’ exon on the 3’ intron/exon border. This event covalently joins the two exons and separates the 3’ exon from the intron, resulting in a free linear intron that then typically circularizes. Group II introns also are self-splicing, and use a similar splicing pathway, except that the initiator of the first reaction is an adenine (A) near the 3’ end of the intron. As with group I introns, there are two transesterification reactions. In the first, the nucleophilic attack by the 2’ hydroxyl on the A within the intron that separates the 5’ exon from the intron, and the intron forms a lariat link with bonds to the 2’, 3’, and 5’ carbons. The second reaction is a nucleophilic attack similar to that for group I introns, which results in joined exons and a free lariat. Group III introns (not shown) are a subtype of group II introns. Twintrons (not shown) are introns within introns and have been described for groups I, II, and III. The interior intron is removed prior to the removal of the exterior intron resulting in splicing of the surrounding exons. Spliceosomal introns undergo the same reactions as those for group II introns, but they are held in their reactive form by spliceosomes, which are small ribonuclear protein structures [43]. Introners are a class of spliceosomal intron and splice in a similar manner. Some of the small RNAs in spliceosomes are similar to those in the longer group II introns, and at least one of the proteins has similarities to ribosomal proteins. Archaeal introns are first cut in two bulges at the splice sites by an endoribonuclease, and an RNA ligase joins the ends of the exons as well as ligating the ends of the exon to form a circular RNA [44]. Similar reactions occur when the large and small rRNA subunits are cleaved from a precursor RNA during maturation of the rRNAs. Some group I introns among Archaea also splice using a bulge-helix-bulge mechanism.

Introns in groups I and II are found in Bacteria and Archaea, and Archaea have an additional “archaeal” type of intron, many of which are also found in tRNA genes in nuclei, mitochondria, and plastids. Whereas the number of introns is limited in prokaryotes, their number and diversity have expanded greatly in eukaryotes, including multiple introns within individual genes and the use of alternative splicing, increasing the sizes of genomes and proteomes [9]. This expansion has led to the expansion of protein diversity without requiring the addition of new genes.

Eukaryotic genomes contain not only group I, group II, and archaeal introns, but also introns classified as spliceosomal, introners, group III, and twintrons. Spliceosomal introns (requiring spliceosomes), introners and group III introns are related to group II introns [6,10], although some reports place introners as diverging from DNA transposons [11–13]. Twintrons are introns within other introns and have been documented for groups I, II, and III [14]. Introns have not only been beneficial in expanding gene and protein diversity, but have added flexibility in gene expression. Mobile introns can move (or move a copy) from one genomic location to another, utilizing either a homing endonuclease, a reverse transcriptase, or both enzymes. Introns may also move about the genome via recombination. MGEs classified as “transposons” can use similar processes to move within a genome, suggesting a common ancestry with introns.

Mobile group I introns insert into new genomic sites using a homing endonuclease to cut a segment of DNA recognized by this enzyme and transfer a DNA copy of the intron to a new site (Fig 1). A DNA transposon is another type of MGE that utilizes a similar mechanism to move into new sites. A mobile group II intron first uses an endonuclease to create a double-stranded break in the DNA and then inserts an RNA version into the DNA, followed by reverse transcription and DNA polymerization to create a copy in the DNA. This process is analogous to insertion of a retrotransposon. In both transposition processes, direct repeats (DRs) result from the insertion [15]. In addition, double-stranded breaks caused by other events can lead to insertion of mobile elements by error-prone mechanisms [16]. Archaeal introns employ a splicing mechanism similar to that used by some group I introns [17] and are integrated into many genes using an endonuclease similar to those used by group I introns. Therefore, archaeal introns also appear to integrate into genomic locations via mechanisms analogous to those used by other introns and transposons.

Here, we present additional evidence supporting intron-transposon connections by analyses of DRs at or near the borders of introns that are found in diverse groups of organisms. These DRs are the hallmark of introns and transposons not classified as introns. The data point to a common theme: introns and transposons are part of a large family of MGEs that move in order to increase their numbers at the expense of the host, even if they are occasionally also useful to the host cell.

## Results

DRs within 40 bp of the 5’ and 3’ exon/intron borders were found in each of the 214 introns examined, including group I, group II, group III, spliceosomal, and archaeal introns, as well as group II and III twintrons (Table 1, Table S1). The DR lengths ranged from 3 to 27 bp, with a mean of 9.8 (sd 2.6). The distances to the exon/intron borders ranged from 0 (spanning the border site) to 37 bp, with a mean of 3.5 (sd 3.9). Most DRs were A/T-rich (Table S1). Few had G/C amounts exceeding 50%. While there were slightly fewer DRs (26%) located in the 5’ exons, the percentages for all other locations were close to the same values (33-37%). DRs on the 5’ exon/intron border were most often paired with DRs on the 3’ intron/exon border (19.6% of all introns examined; Fig 2). DRs near the 5’ end of the intron were most often paired with DRs in the 3’ exon (18.2%), while DRs in the 5’ exon were most often paired with DRs near the 3’ end of the intron (14.4%). All of these approximated the total lengths of the respective introns. All other combinations of DRs in the 5’exons, exon/intron borders, and 3’ exons were either shorter or longer than the respective introns (Fig 2). Each of the intron types exhibited a variety of DR arrangements (Fig 2, table on right). The most common DR location for spliceosomal introns (37.8%) was the exon/intron borders; and the most common location for group I, II, and III introns was one DR within the intron and the other in one of the exons, without altering the length of the intron (yellow and black bars in Fig 2). Each of these arrangements maintains the lengths of the mature RNAs after splicing, but two of the three change their end sequences. Intron versions that decrease (orange bars) and increase (red bars) the length of the splice products were also found. Although fewer archaeal introns were analyzed, most would alter the length of the mature RNA.

**Table 1.** Summary of direct repeats at or near the 5’ and 3’ exon/intron borders, including locations, frequencies, lengths, and distances relative to the exon/intron border.

| Intron Type | Cell Location | Taxa Where Present<sup>a</sup> | 5' Repeat Location Frequency |  |  | 3' Repeat Location Frequency |  |  | Mean Repeat Length (bp)<sup>b</sup> | Mean Distance to e/i Border (bp)<sup>c</sup> |
| --- | --- | --- | --- | --- | --- | --- | --- | --- | --- | --- |
|  |  |  | 5' exon | exon/intron border | intron | intron | intron/exon border | 3' exon |  |  |
| Group I | Nuclear | Al,Am, An,Fu, Pt,Rh, St | 2/12 | 6/12 | 4/12 | 0/12 | 6/12 | 6/12 | 6.2 ± 3.4 | 0.0 ± 0.0 (0-0) |
|  | Mitochondrial | Al,Am, An,Ex, Fu,Pt, St | 1/2 | 0/2 | 1/2 | 0/2 | 0/2 | 2/2 | 8.0 ± 3.5 | 3.6 ± 5.3 (0-14) |
|  | Plastid | Pt,St | 2/19 | 5/19 | 12/19 | 4/19 | 5/19 | 10/19 | 9.9 ± 3.6 | 6.6 ± 7.1 (0-21) |
|  | Prokaryote | Ba | 2/2 | 0/2 | 0/2 | 1/2 | 1/2 | 0/2 | 16.0 ± 3.5 | 4.2 ± 4.2 (0-12) |
|  | **TOTAL** | ----- | **7/35** | **11/35** | **17/35** | **5/35** | **12/35** | **18/35** | **8.8 ± 4.2** | **4.7 ± 6.6 (0-21)** |
| Group II | Nuclear | Fu | 0/1 | 1/1 | 0/1 | 0/1 | 1/1 | 0/1 | 10.5 ± 0.5 | 0.0 ± 0.0 (0-0) |
|  | Mitochondrial | Al,An, Ex,Fu, Ha,Pt, Rh, St | 12/35 | 12/35 | 11/35 | 17/35 | 8/35 | 10/35 | 12.0 ± 3.9 | 8.2 ± 9.0 (0-37) |
|  | Plastid | Pt | 19/66 | 17/66 | 30/66 | 24/66 | 20/66 | 22/66 | 11.4 ± 4.6 | 4.8 ± 6.4 (0-31) |
|  | Prokaryote | Ar,Ba | 1/2 | 1/2 | 0/2 | 0/2 | 1/2 | 1/2 | 10.8 ± 2.4 | 8.0 ± 8.0 (0-17) |
|  | **TOTAL** | ----- | **32/104** | **31/104** | **41/104** | **41/104** | **30/104** | **33/104** | **11.6 ± 4.3** | **5.9 ± 7.6 (0-37)** |
| Group III | Plastid | Ex | 7/25 | 10/25 | 8/25 | 11/25 | 6/25 | 8/25 | 13.8 ± 5.1 | 5.0 ± 5.5 (0-22) |
| Spliceosomal | Nuclear | Al,Am, An,Ex, Fu,Pt, Rh,St | 6/45 | 26/45 | 13/45 | 16/45 | 21/45 | 9/45 | 11.1 ± 5.1 | 3.6 ± 5.9 (0-29) |
| Archaeal | Nuclear | Al,An, Fu,Pt,St | 1/1 | 0/1 | 0/1 | 0/1 | 0/1 | 1/1 | 5.5 ± 0.5 | 6.5 ± 6.5 (0-13) |
|  | Prokaryote | Ar | 2/4 | 2/4 | 0/4 | 1/4 | 2/4 | 0/4 | 10.9 ± 1.5 | 2.8 ± 2.4 (0-6) |
|  | **TOTAL** | ----- | **3/5** | **2/5** | **0/5** | **1/5** | **2/5** | **1/5** | **9.8 ± 2.6** | **3.5 ± 3.9 (0-13)** |
| **Grand Totals** |  |  | **55/214 (26%)** | **80/214 (37%)** | **79/214 (37%)** | **74/214 (35%)** | **71/214 (33%)** | **69/214 (32%)** | **10.5 ± 2.4** | **4.4 ± 2.6** |

<sup>a</sup> Taxon names: Al = Alveolata; Am = Amoeboae; An = Animals; Ar = Archaea; Ba = Bacteria; Ex = Excavata; Fu = Fungi; Ha = Hacrobia; Pt = Plants; Rh = Rhizaria; Sr = Stramenopiles. Presence of intron types from Rogers<sup>19</sup>.
<sup>b</sup> Mean and standard deviation are provided.
<sup>c</sup> Mean and standard deviation are provided. Range of lengths provided in parentheses.

**Figure 2.**
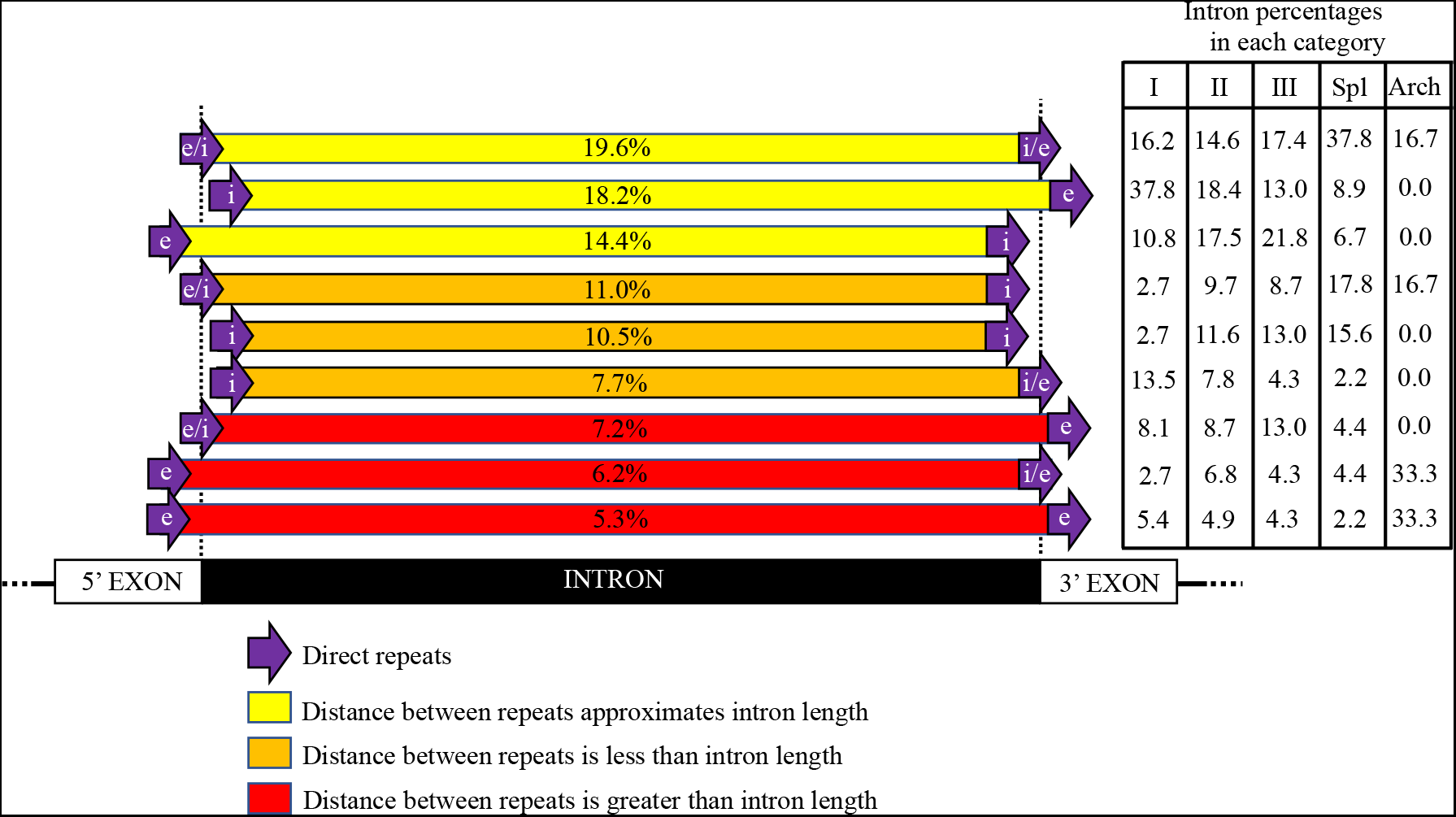
Frequencies of distances between the DRs in the vicinity of the respective introns, arranged in decreasing frequencies. In the most frequent arrangement (19.6% of the cases), the 5’ and 3’ DRs spanned the exon/intron borders. The yellow bars indicate DRs separated by the length of the intron plus the several nucleotides added to create the DR (52.2% of the cases). The orange bars indicate DRs separated by less than the length of the intron (29.2% of the cases); the red bars in indicate DRs separated by more than the length of the intron (18.7% of the cases). The table shows the frequencies (as percentages) for each of the DR-pair categories for each type of intron.

In summary, each of the introns analyzed was bounded by a pair of short DRs. For all introns analyzed, the locations of some of the DRs indicated that a spliced RNA transcript would rectify the gene defect caused by intron insertion. For each category of intron, however, most DR locations would alter the length and/or end sequences of the mature RNA potentially leading to a defective gene product.

## Discussion

Previously, we documented the locations of numerous DRs caused by insertion of transposons within the rDNA loci of the eukaryotic nucleus [15]. Here, we report DRs at or adjacent to the borders of every intron examined in seven major taxa of Eukarya, Bacteria, and Archaea, and in mitochondria and plastids (Table 1, Fig 2, Table S1). The most plausible conclusion is that transposons and introns are related to the extent that they represent members of a large family of MGEs. Mechanistic and evolutionary aspects of this relationship, however, have been obscure. Mobile group I introns carry at least one gene that is an endonuclease, analogous to the endonucleases present in autonomous DNA transposons (Figs 1 and 3). Integration occurs at the break site caused by the endonuclease. The process produces DRs at the border sites. Integration can also occur at other double-strand breaks not caused by an endonuclease, using an error-prone repair system [15,16]. Similarly, mobile group II introns have at least one gene that is a reverse transcriptase related to the reverse transcriptase present in retrotransposons (Figs 1 and 3). These introns integrate at double-strand break sites, some of which are caused by endonucleases, and this integration also produces DRs at the borders of the inserted mobile element (Figs 2 and 3). The integration of a DNA element can, presumably, occur at any double-strand break point. Autonomous transposons carry the genes necessary for their movement into new sites, while nonautonomous versions can still move if autonomous versions are present to supply the necessary gene products for transposition. This is analogous to the relationship between mobile and non-mobile introns that are still capable of movement.

**Figure 3.**
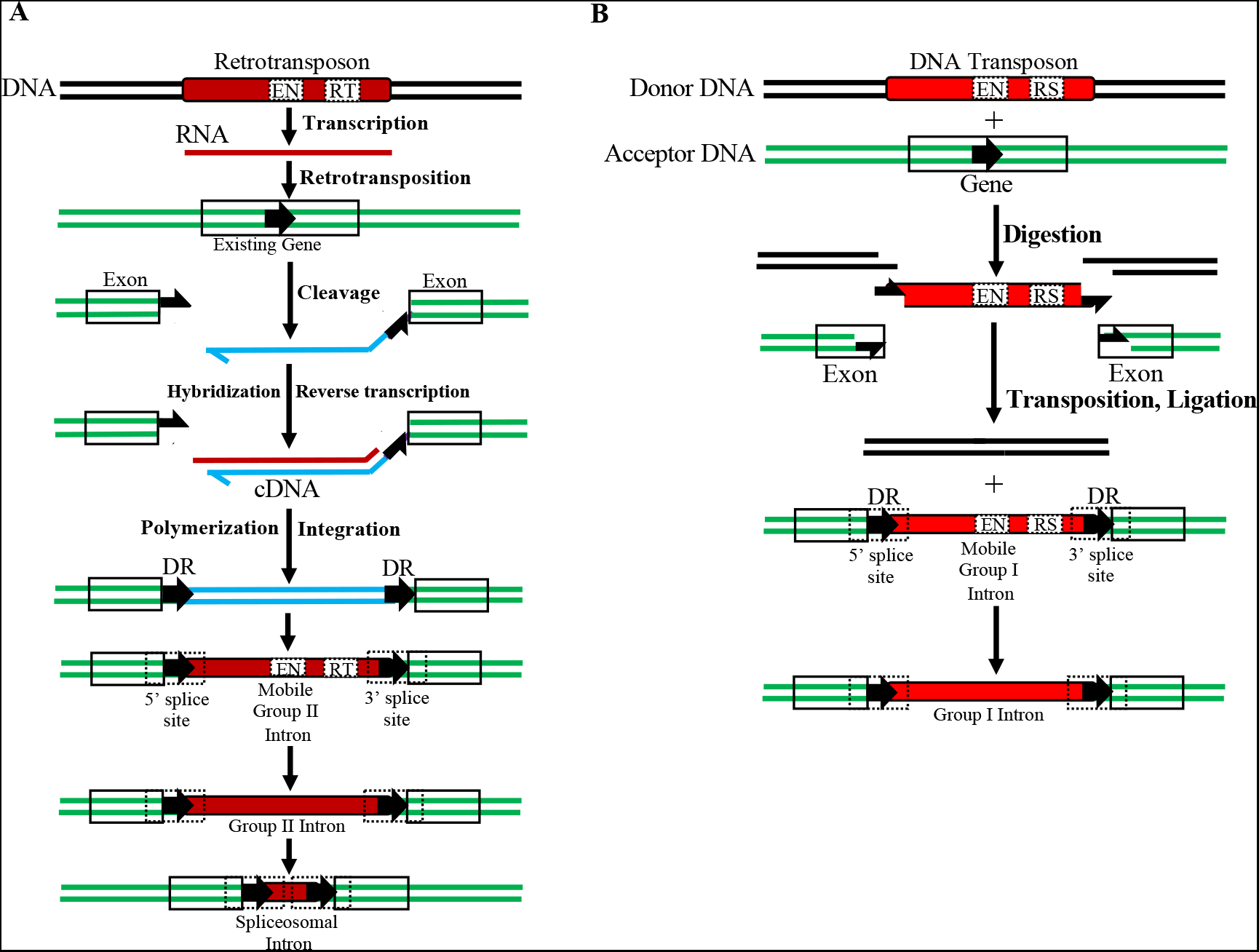
Proposed mechanisms for conversion of transposons into introns. Insertions follow the general pathways for transposition. (A) Conversion of a retrotransposon into a mobile group II intron. Both encode a reverse transcriptase (RT), and some also encode an endonuclease (EN). DRs are produced in the process, although the 5’ and 3’ splice sites appear either within the DRs or close to the DRs within the exons and intron (dashed boxes; see Table 1 and Fig 2). The mobile group II introns may then change further, becoming introners or nonautonomous group II, group III, and spliceosomal introns. Although the length of spliceosomal intron shown here is short, these introns can vary from tens to tens of thousands of nucleotides. (B) Origin of mobile group I introns from DNA transposons, both of which often encode an EN and a resolvase (RS) and/or other genes. Mobile group I introns then led to group I introns without internal genes, including introns from 60 to thousands of nucleotides in length. Some introns produce small RNAs that may control gene and/or RNA expression. Twintrons are introns within other introns and have been described for group I, II, and III introns. Mobile archaeal introns contain coding regions for endonucleases similar to those in group I introns, so their insertion steps may be similar to those for group I introns.

### Commonalities between Transposons and Introns

Similarities between transposons and introns have long been recognized. Rogozin et al. [18] reported similar properties for the reverse transcriptases used by retrotransposons and group II introns (and retroviruses), and Chalamcharla et al. [10] described the evolution progression from group II introns to spliceosomal introns. This progression is also illustrated by the relative numbers of group II and spliceosomal introns in rDNA. Group II introns are found in older portions of rDNA and then decline in more-recently added sections. Their decline coincides with increases in the number of spliceosomal introns in both the large subunit (LSU) and small subunit (SSU) genes [19]. Introners appear to be a type of spliceosomal intron capable of rapid spread through genomes either by reverse splicing or transposition [5,11–13,20]. Although the link between DNA transposons and group I introns is more tenuous, both use endonucleases that target specific “homing” sites [21]. Archaeal introns splice by a different mechanism than do most group I introns (Fig 1), but both use a similar endonuclease to initiate integration into the DNA. Bioinformatic analyses have indicated that within Archaea, archaeal introns were derived from group I introns, including some group I introns that splice via a bulge-helix-bulge mechanism that is the hallmark of archaeal splicing [17]. Rogers [19] concluded that group I introns appeared in the earliest-evolving segments of the SSU and LSU rRNA genes, while archaeal introns appeared in subsequently-added segments of those genes, and that group I introns are found exclusively in the oldest portions of the SSU rRNA genes [19].

Introns and transposons are found in all types of genes (mRNA, tRNA, sRNA, and rRNA) and in all groups of eukaryotes. Additionally, the cleaving of SSU and LSU rRNAs from a long precursor molecule is similar to splicing of archaeal introns, representing a link between archaeal introns and rDNA internal transcribed spacers (in bacteria, archaea, and eukaryote ITS1 and ITS2; [22]). Whereas rDNA was proposed as a source of introns migrating elsewhere in the genome, the immigration of introns and transposons into rDNA may have created new sections in rRNAs [19]. Furthermore, the high copy numbers of rDNA units present large targets for incoming MGEs, as well as a resource for mobilization to other loci. Introns in rDNA are concentrated in the older parts of the SSU and LSU rRNA genes and are rare in the more recent additions to the rDNA, suggesting an early emergence of introns, followed by a reduction in the rate of intron spreading [19].

In conclusion, although transposons and introns are typically classified as distinct categories of MGEs, both use similar (or the same) enzymatic agents to achieve the same selfish result: proliferation and spread of themselves. The host cells later exploited variant transposons and introns for their own benefit (alternative splicing, for example).

### Conversion of Transposons into Introns

Our proposed mechanism for transposon-to-intron conversion is analogous to transposition alone, with the addition of at least two splice sites (Figs 2 and 3). The arriving DNA element may insert in a location that already includes borders compatible with splicing or the element may have a sequence resembling an exon/intron border [23], while the other border already exists in the flanking sequences. Integration itself may create new splice sites comprising nucleotides from the exon and the new intron [12]. Cryptic splice sites can alter the mature RNAs, and splicing errors are common [24,25]. Our results point to a chaotic process in which an MGE is inserted into a gene, followed by heavy selection favoring spliced products that are still functional. This causes splice sites to be selected that may or may not coincide with the length of the inserted MGE. The insertion of an intron (or transposon) may thus create alternative RNAs that affect species fitness [26–28]. Most insertions will be detrimental, but occasionally a beneficial intron may emerge. In some cases introns have become intertwined, creating a twintron [14], and introns themselves are hotspots for additional transposon integration [29].

Introns exhibit plasticity. In one case, a group I intron near the 3’ end of the SSU rRNA gene in a fungus folds in both a group I as well as a group II form, with intron borders shifted by only two nucleotides [19,30,31]. In another case, several species of fungi contain group I introns within the SSU rDNA that range from 62-78 nucleotides. Although the entire canonical central portions of the introns are missing, part of the adjacent 3’ SSU and ITS1 of the rDNA compensate for the missing sections to allow accurate splicing of the SSU rRNA [19,32,33]. These findings indicate that exon/intron lengths and borders can change substantially.

Whereas the yellow bars in Figure 2 indicate that intron-length inserts comprise most (52.2%) of all inserts analyzed, the fraction of all introns that carry DRs at both borders of the original intron was highest for spliceosomal introns (37.8%) among the five intron categories. This relatively low flexibility in DR placement probably reflects the need to produce a functional protein, whereas for the other four categories splicing produces mature transcripts more likely to be functionally compromised. In all cases, however, most of the transcripts do not represent a “clean” removal of intronic RNA to rescue the product of the original gene.

Most DR arrangements for group I introns included one DR near the 5’ end of the intron and the other DR near the 5’ end of the 3’ exon, without altering intron length. Group I introns were almost exclusively found in rRNA and tRNA genes. Because there are no constraints to maintain a reading frame for these RNAs, selection appears to be primarily based on whether the mature RNAs can fold into their functional conformations. The DR pairs flanking group II and III introns (found primarily in organelles and prokaryotes) are less positionally constrained than either group I or spliceosomal introns, but the distance between DRs still approximates the intron lengths. Most archaeal introns examined had DR distances that would alter one or both exons. Because most tRNA-gene introns are in the anticodon loop segment, some variability in the mature tRNAs and their anticodons should result. The evolution of tRNAs and their cognate aminoacyl tRNA transferases (aaRSs) is complex, including some aaRSs that charge biochemically-similar tRNAs and modify tRNAs leading to a diverse set of tRNAs [19,34]. Changes in tRNA specificity caused by intron insertion may, therefore, be tolerated or even occasionally beneficial during evolution.

In 29.2% of the cases the distance between the DRs was shorter than the intron (Fig 2, orange bars), creating an extension of one or both exons and mature RNAs that would be longer than the ancestral version before intron insertion. Conversely, in 18.7% of the cases (Fig 2, red bars) the distance between the DRs exceeded the intron length, which would shorten the mature RNA. Lengthened and shortened RNAs should negatively affect the resulting gene products, illustrating the deleterious effects of most introns on the cell. When the host cell does occasionally survive an intron insertion, however, the intron itself is the primary beneficiary of the insertion event.

### Data summary and interpretation

We have identified DRs at or near the splice sites for 214 diverse introns and find their properties analogous to those of DRs surrounding transposons. Based on those DRs, genes within autonomous transposons and mobile introns, and the mechanisms of movement of transposons and mobile introns, we infer the following (Fig 4): Introns likely originated from transposons; Introns in groups II and III were derived from retrotransposons; Spliceosomal introns are derivatives of group II introns; Introners are derivatives of group II introns, having many characteristics of group II introns (Table S2), but have adapted to spread rapidly through populations; Group I introns originated from DNA transposons; Archaeal introns were derived from group I introns. All types of transposons and introns exist in autonomous and nonautonomous versions (black and red fonts, respectively, in Fig 4)

**Figure 4.**
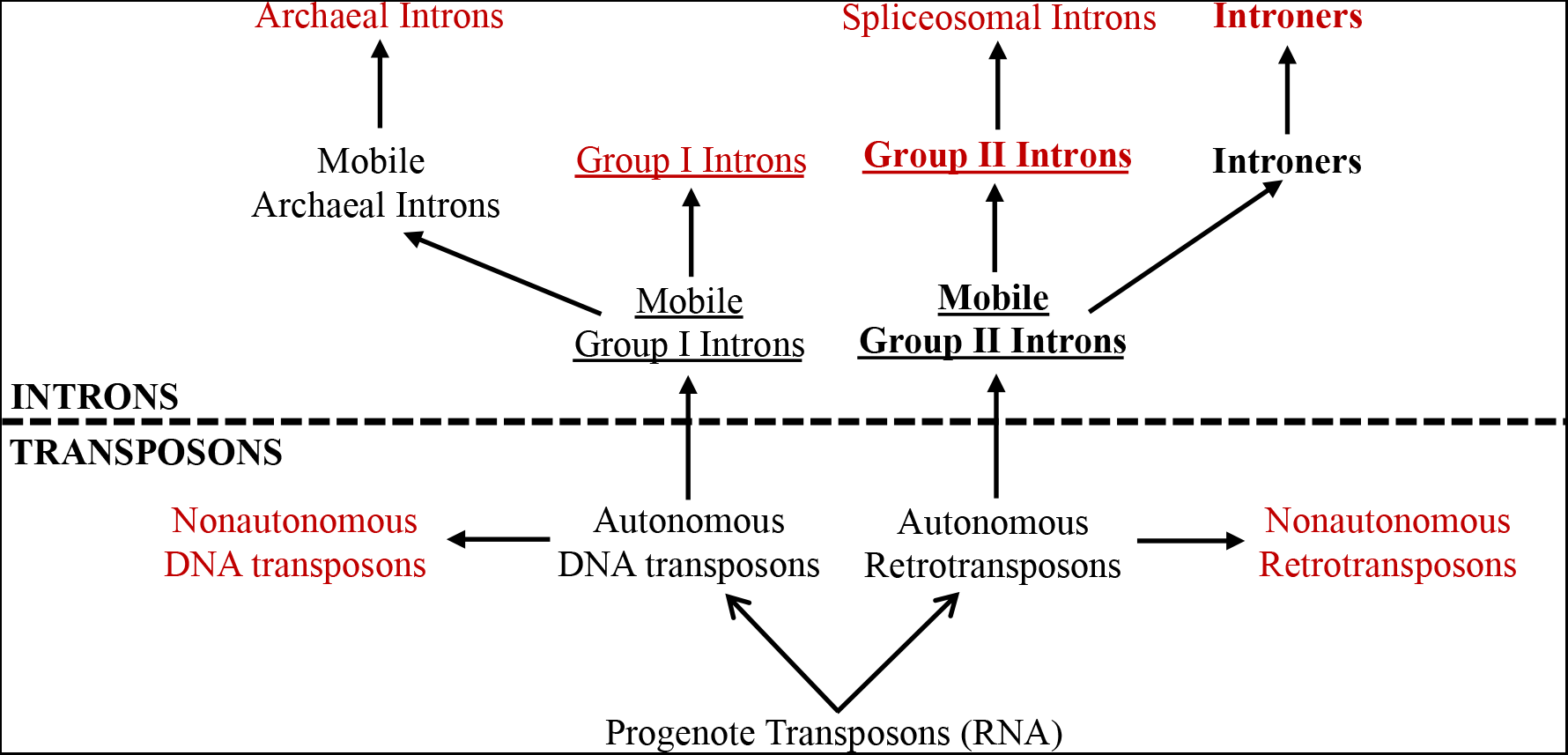
Proposed evolutionary relationships among transposons and introns. Transposons likely began as autonomous RNA molecules that could invade other nucleic acids. These diversified into autonomous DNA transposons, which contained a gene for an endonuclease, and autonomous retrotransposons that carried a reverse transcriptase gene. They often carried other genes that aided transposition. Members of each class lost mobility genes that then were supplied by mobile versions. Some members of each autonomous version, mobile group I and group II introns, could also splice themselves out of the RNA versions which was an advantage because integration into genes would often damage the RNAs. Again, mobility genes were lost and had to be provided by autonomous versions. Derivatives of one type of group I became archaeal introns. Derivatives of group II introns became spliceosomal introns and introners that splice similarly (Fig 1; [5,11]). Most introners lack terminal inverted repeats (Table S2) and are therefore placed within the retrotransposon lineage. Both transposons and introns are “selfish” elements that are autonomous (black font) or nonautonomous (red font). Underline = self-splicing introns; bold font = capable of reverse splicing.

### Concluding remarks

Why genes in pieces? This question has existed since introns were discovered [35–37]. Most proposed answers emphasize the benefits to organisms carrying introns, such as expanding the proteome and increased flexibility of gene expression. Although these organismal benefits are unquestioned, we suggest that the original beneficiary was the intron DNA itself acting as a selfish MGE, with organismal benefits appearing later in evolution. Damage to DNA in all genomes is unavoidable. Repair of the most severe type of damage, a double-strand break (DSB), must be successful if the cell is to survive. We previously proposed that the insertion of a transposon involves a DSB requiring repair [15], and we now propose the same for insertion of an intron. In bacteria, the fraction of the genome occupied by transposons plus transposon-defense genes is only about one or two percent [38–40] and an even smaller fraction is attributed to introns. The fraction in eukaryotic nuclei is larger, the genomes are larger, and the number of introns is in the thousands and tens of thousands [41]. In many eukaryotes the number of nucleotides in introns of a gene is greater than that in the exons, and splicing errors are common [42]. The major unanswered question is what accounts for the permissive nature of eukaryotic genomes that has allowed the dramatic rise of introns, transposons, and other MGEs?

## Materials and Methods

All sequences including introns were retrieved from NCBI during July through September 2022. Seven major taxonomic groups were sampled: Alveolata [(*Symbidiodinium microadraticum* — dinoflagellate symbiont of jellyfish, nuclear spliceosomal introns, accession number LSRX01000131)]; Animalia [(*Homo sapiens*, nuclear dystrophin gene spliceosomal introns, accession numbers AJ27220 and U60892), (*Ricordea yuma* — coral, mitochondrial group I intron, accession number H308005), (*Carcharhinus leucas* — bullshark, nuclear rDNA ITS group I intron, accession number JN039366), (*Isurus oxyrhinchus* — shortfin mako shark, nuclear rDNA ITS group I intron, accession number MK079238), (*Stegobium paniceum* — bread beetle, nuclear rDNA group I introns, accession number D49657)]; Archaea [(*Methanospirillum hungatei*, tRNA archaeal intron, accession number NC_000916), (*Staphylothermus marinus*, rRNA archaeal intron, accession number NR_076485), *(Thermofilum pendens*, tRNA archaeal introns, accession number NZ_AASJ01000001)], Archaeplastida [(*Magnolia macrophylla* — bigleaf magnolia, plastid group I and group II introns, accession number AY687352), (*Triticum aestivum* — wheat, mitochondrial group II introns, accession numbers X75036, X57164; nuclear spliceosomal introns, accession number AJ512822; plastid group I and group II introns, accession number NC_002762), (*Vicia faba* — broad bean, mitochondrial group I and group II introns, accession number KC189947; nuclear spliceosomal introns, accession numbers AM886054, AJ277286; plastid group I and group II introns, accession number MT120813)], Bacteria [(*Bacillus* sp. BSG40, bacterial group I intron, accession number AJ309312), (*Streptococcus agalactiae*, bacterial group II introns, accession numbers AF494487, AY189967), (*Thermotoga subterranea*, bacterial group I intron, accession number AJ556793)], Excavata [(uncultured trypanosome, nuclear spliced leader spliceosomal intron, accession number KR056281), (*Euglena gracilis*, plastid group II and group III introns, including some twintrons, accession number Z11874)], Fungi [(*Amplexidiscus fenestrafer*, mitochondrial group II intron, accession number MH308002), (*Phialophora verrucosa*, nuclear rDNA group I intron, accession number JF414781), (*Pencillium oblatum*, nuclear rDNA group I intron, accession number AB033529), (*Cenococcum geophyllum*, nuclear rDNA group I and group II introns, accession number Z11998), (*Xylaria polymorpha*, nuclear rDNA group I intron, accession number AB01404), (*Saccharomycodes ludwigii*, nuclear archaeal intron, accession number NC_060204), (*Mycoarachis inversa*, nuclear rDNA spliceosomal introns, accession number AB012953), (*Saccharomyces cerevisiae*, nuclear spliceosomal and group I introns, accession number LBMA01000012), (*Schizosaccharomyces pombe*, mitochondrial group II introns, accession number NC_001326), (*Scytalidium dimidatum*, nuclear rDNA group I intron, accession number AF258603)]. The 5’ and 3’ intron borders that were identified in the NCBI sequences were used as the starting points for searches of DRs of at least 3 bp, and within 40 bp from the borders. The longest identical DRs were retained. Identical DRs were recorded (e.g., in *Stegobium paniceum* rDNA group I (2), aacggg/aacggg; Table S1), as well as many that were highly similar containing 1-2 bp differences for short repeats (e.g., *H. sapiens* dystrophin gene spliceosomal intron, ttttg/ttttag; Table S1) up to 8 bp differences for long repeats (e.g., *Magnolia macrophylla*, plastid rpoC1 gene group II intron, tatccaaagctaccctagggg/tattaaagctagtttagtgtg).

## Acknowledgments

We thank George Miklos and Luca Comai for their useful comments.

## Author Contributions

Planning and conceptualization, AJB and SOR; Bioinformatic analyses, SOR; Writing and editing manuscript, AJB and SOR; Figures, SOR.

## Supporting Information

**Table S1.**
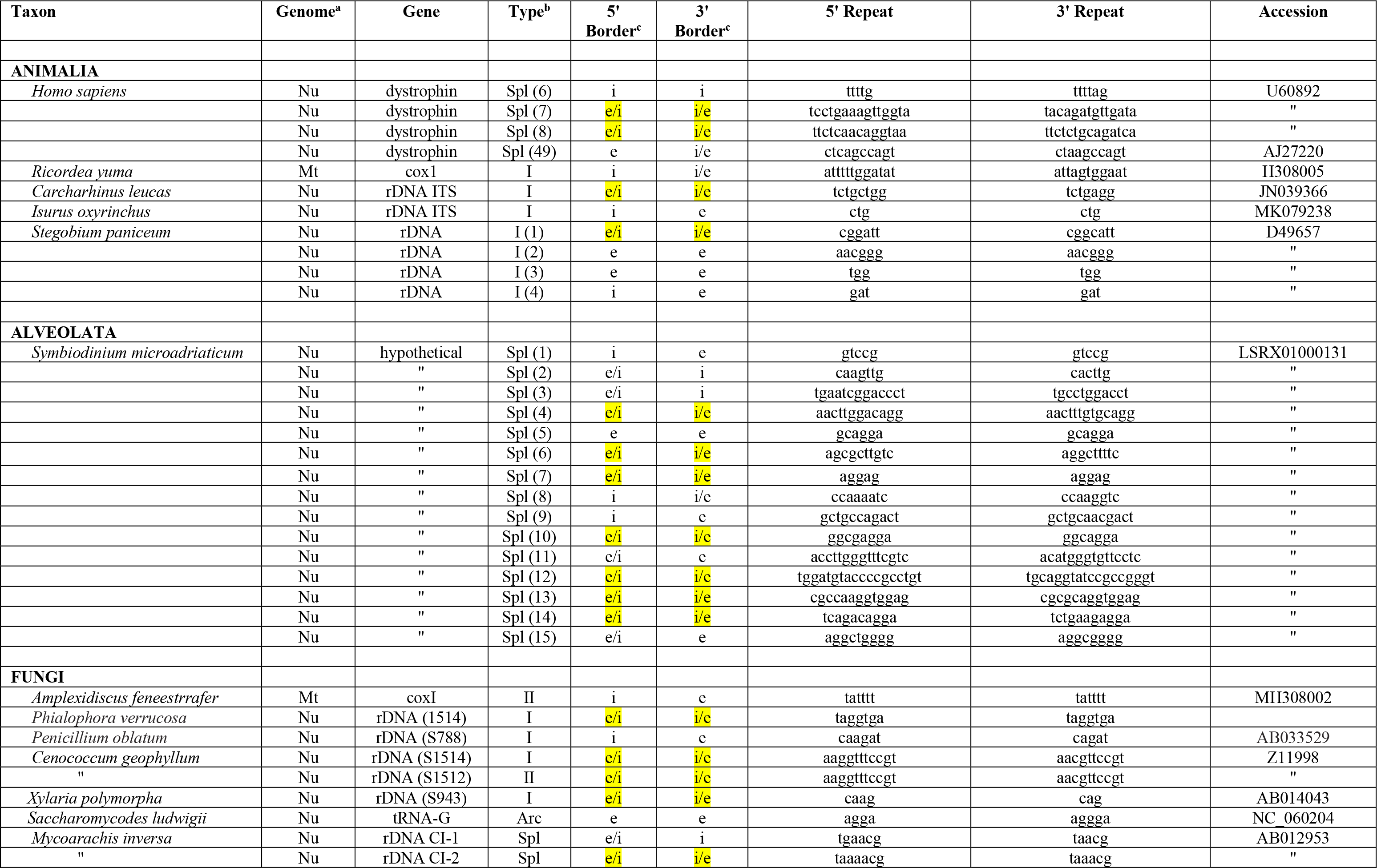
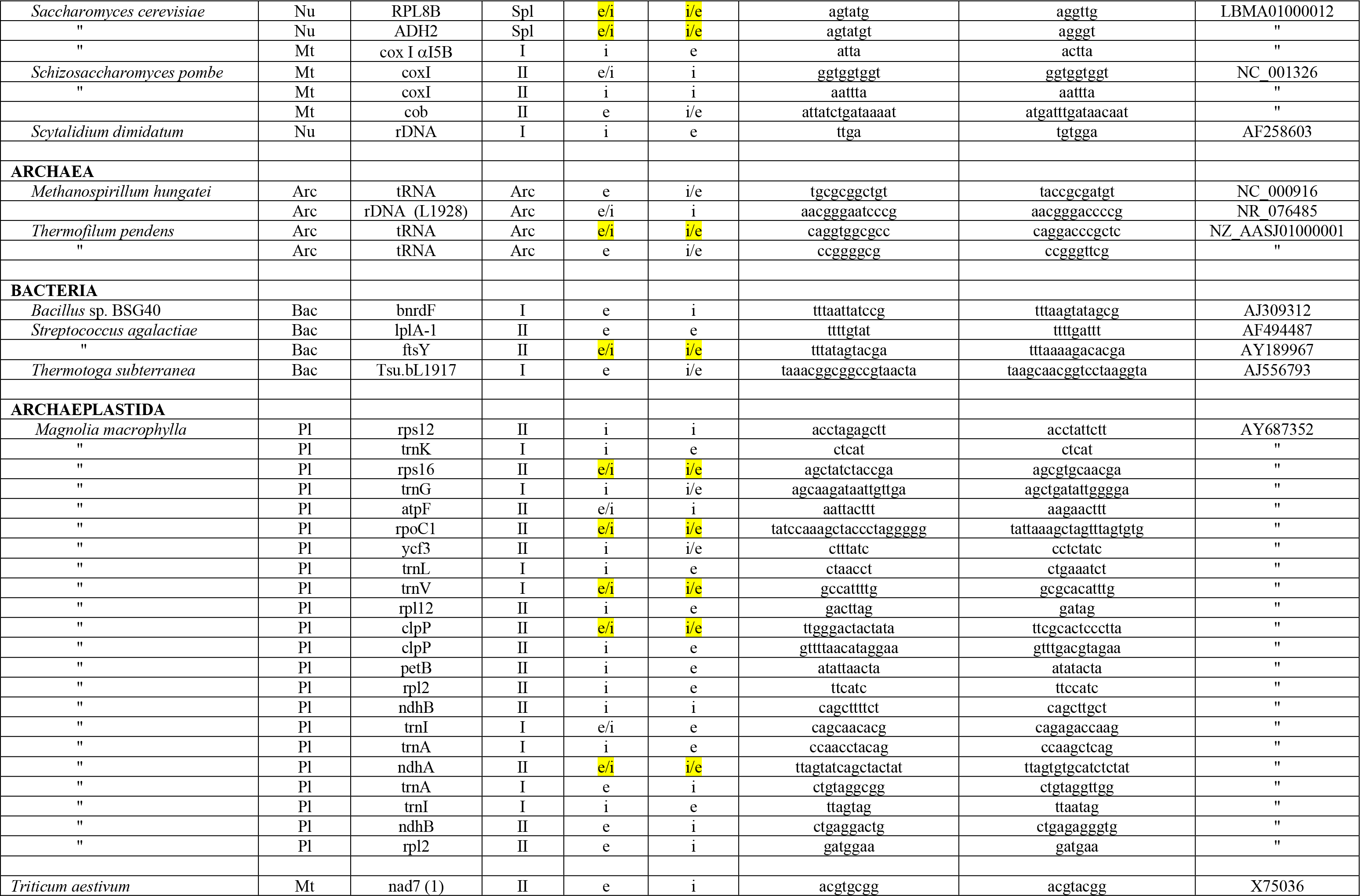
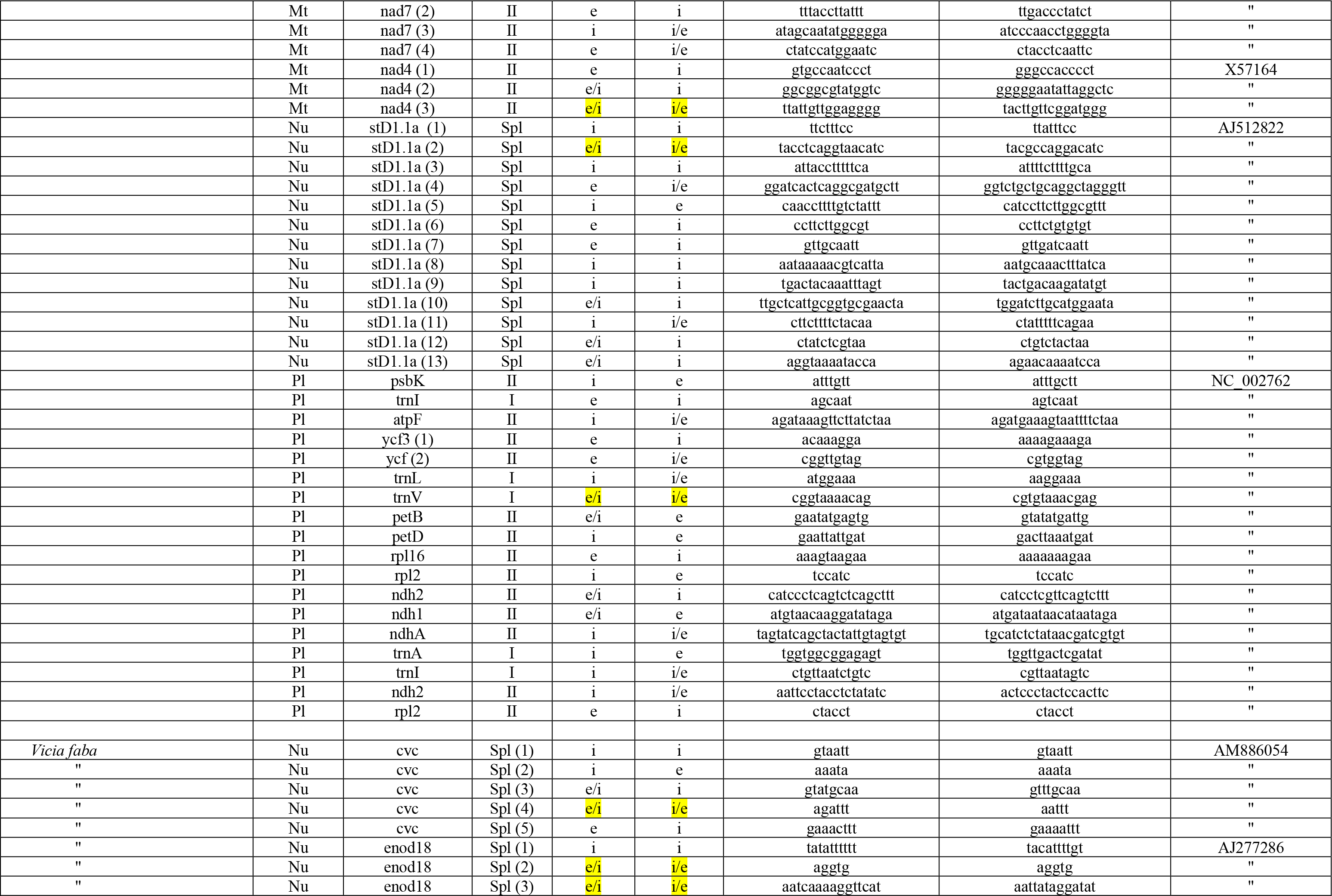
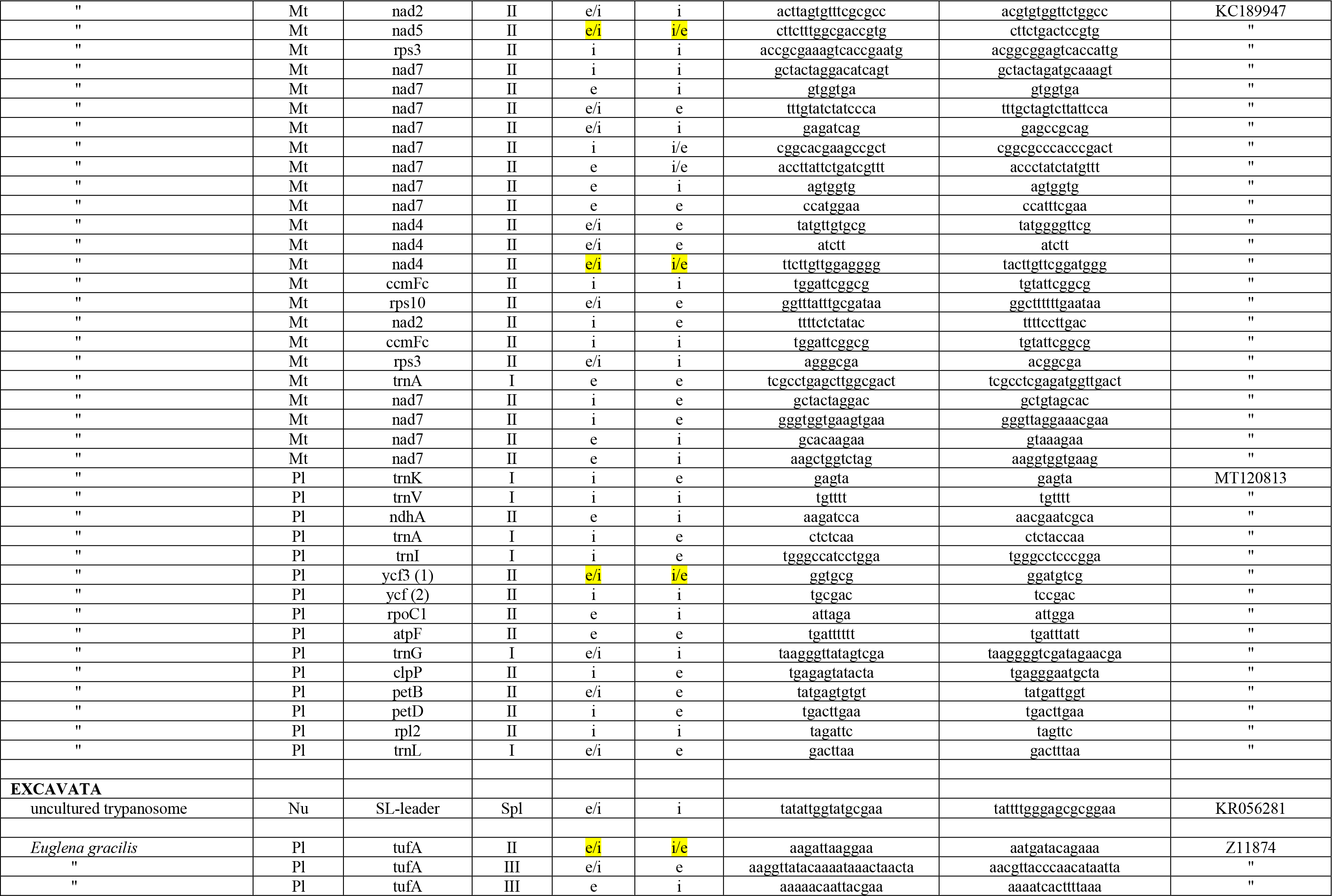
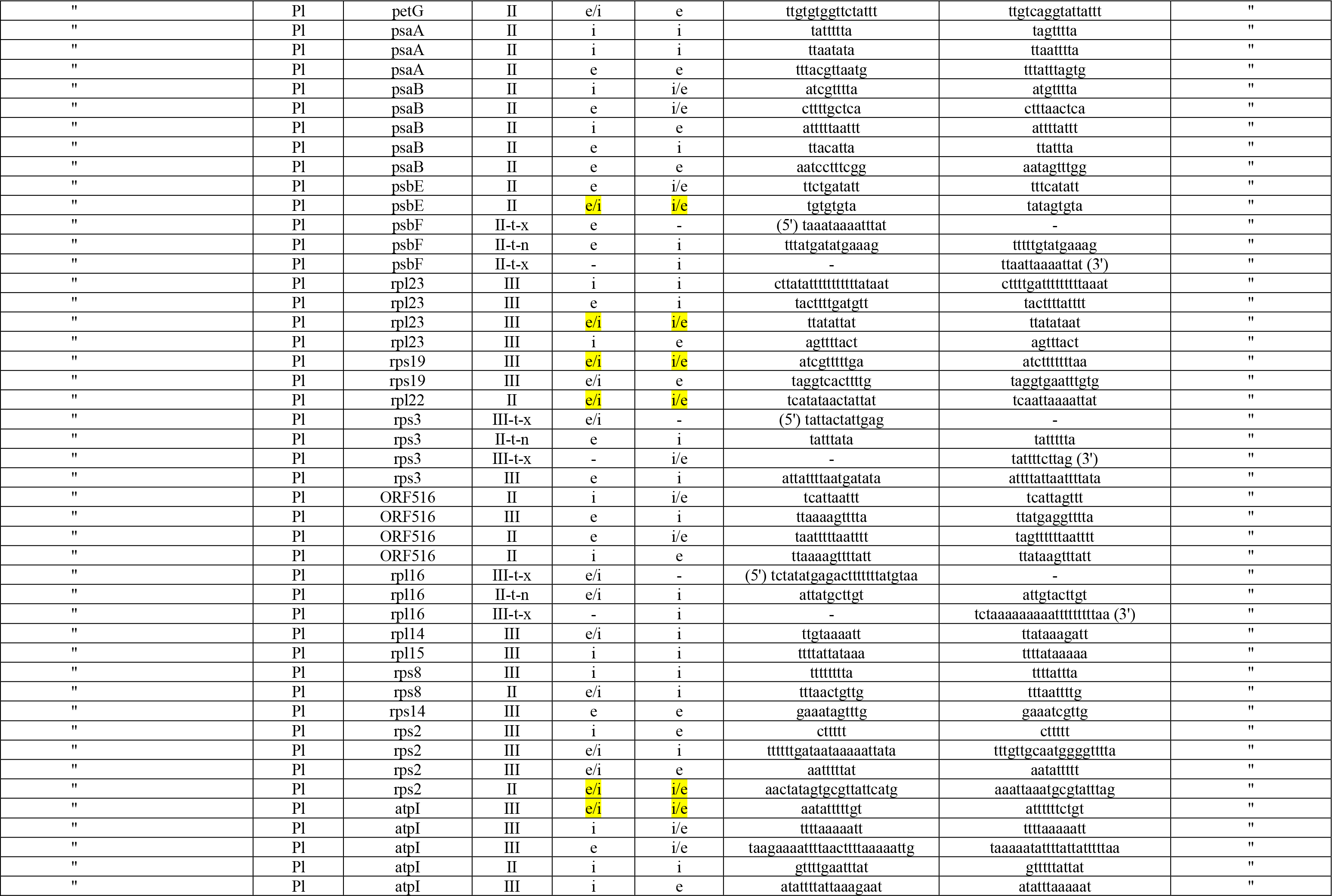
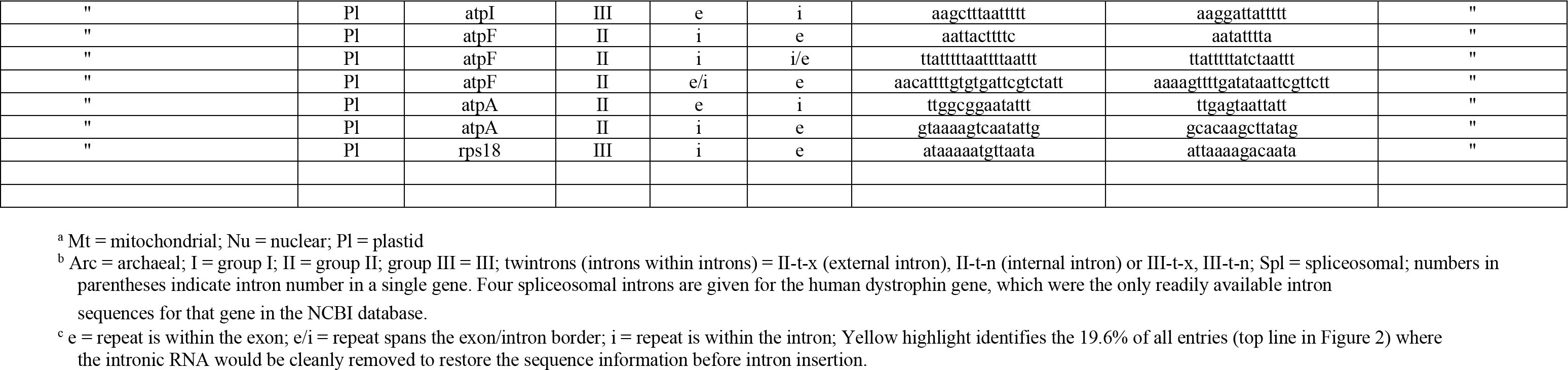
Direct Repeats at or adjacent to the 5’ and 3’ exon/intron borders.

**Table S2.**
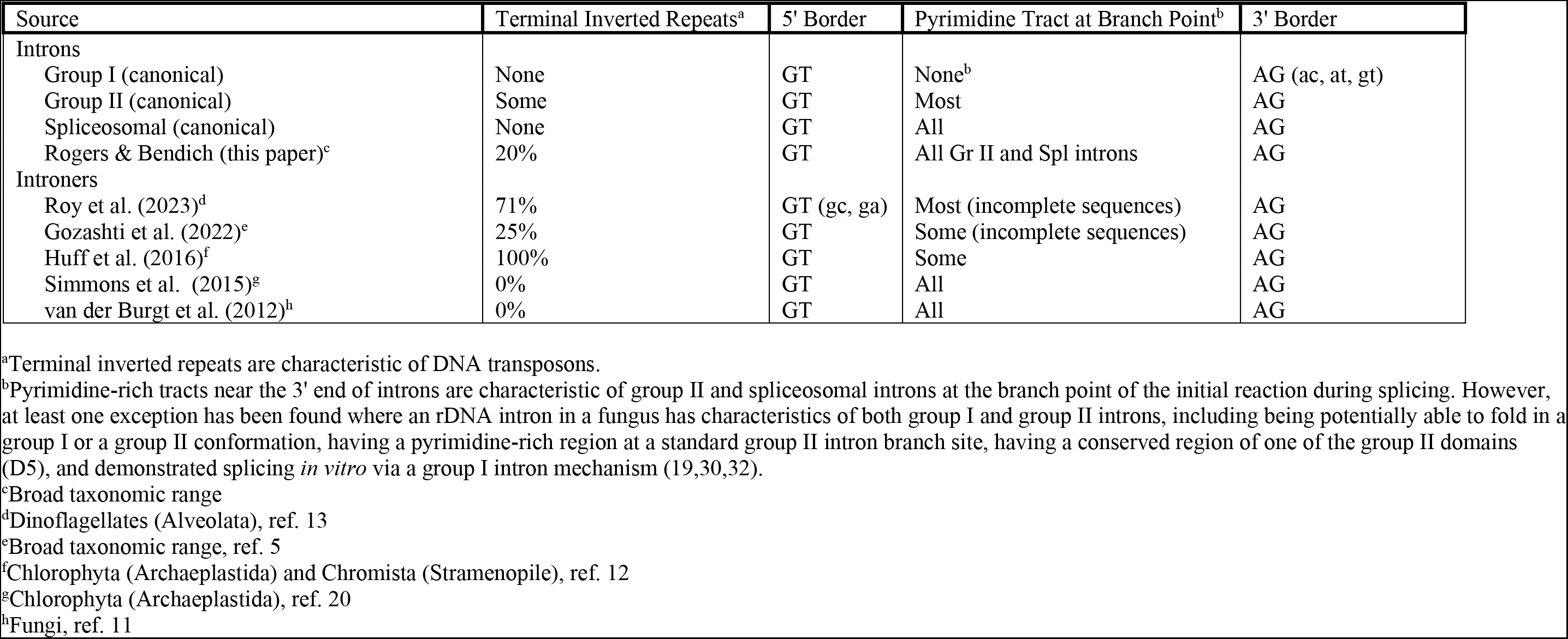
Comparisons of intron and introners Most introns and introners have well-defined borders, usually GT at the 5’ exon/intron border and AG at the 3’ border, although some alternative nucleotides have been reported (as noted in lower case font). Most introners have pyrimidine-rich tracts within a region that suggests a group II (retrotransposon) mode of splicing. Some have terminal inverted repeats that are similar to those found among DNA transposons. However, the terminal inverted repeats are limited to few taxa.

## References

1. Jo B-S, Choi SS. Introns: the functional benefits of introns in genomes. Genomics Inform. 2015;13:112–118. doi: 10.5808/GI.2015.13.4.112.

2. Perotto S, Nepote-Fus P, Saletta L, Bandi C, Young JPW. A diverse population of introns in the nuclear ribosomal genes of ericoid mycorrhizal fungi includes elements with sequence similarity to endonuclease-coding genes. Mol. Biol. Evol. 2000;17:44–59.

3. Randau L, Söll D. Transfer RNA genes in pieces. EMBO Rep. 2008;9:623–628. doi: 10.1038/embor.2008.101.

4. Shivji M, Rogers SO, Stanhope M. Molecular studies on sharks. *Shark Tagger* 1995 Summary, NOAA. 1966;p.15.

5. Gozashti L, Roy SW, Thornlow B, Krameer A, Ares M Jr, Corbett-Detig R. Transposable elements drive intron gain in diverse eukaryotes. Proc. Natl. Acad. Sci. U.S.A. 2022;119(48):e2209766119. doi.org/10.1073/pnas.2209766119.

6. Novikova O, Belfort M. Mobile group II introns as ancestral eukaryotic elements. Trends in Genet. 2017;33:773–783. doi.org/10.1016/j.tig.2017.07.009.

7. Roy SW. The origin of recent introns: transposons? Genome Biol. 2004;5:251. http://genomebiology.com/2004/5/12/251.

8. Yenerall P, Zhou L. Identifying the mechanisms of intron gain: progress and trends. Biol. Dir. 2012;7:29.

9. Kumari A, Sedehizadeh S, Brook JD, Kozlowski P, Wojciechowska M, Differential fates of introns in gene expression due to global alternative splicing. Human Genet. 2022;141:31–47. doi: 10.1007/s00439-021-02409-6.

10. Chalamcharla VR, Curcio MJ, Belfort M. Nuclear expression of a group II intron is consistent with spliceosomal intron ancestry. Genes Devel. 2010;24:827–836. doi: 10.1101/gad.1905010.

11. van der Burgt A, Severing E, de Wit PJGM, Collemare J. Birth of the new spliceosomal introns in fungi by multiplication of introner-like elements. Curr. Biol. 2022;22:1260–1265. doi:10.1016/j.cub.2012.05.011.

12. Huff JT, Zilberman D, Roy SW. Mechanism for DNA transposons to generate introns on genomic scales. Nature 2016;538:533–536. doi: 10.1038/nature20110.

13. Roy SW, Gozashti L, Bowser BA, Weinstein BN, Larue GE, Corbett-Detig R. Intron-rich dinoflagellate genomes driven by introner transposable elements of unprecedented diversity. Curr. Biol. 2023;33:189–196. doi.org/10.1016/j.cub.2022.11.046.

14. Hafez M, Hausner G. Convergent evolution of twintron-like configurations: One is never enough. RNA Biol. 2015;12:1275–1288.

15. Bendich AJ, Rogers SO. Ribosomal spacers are filled with transposon remnants. bioRxiv 2023. doi.org/10.1101/2023.02.20.529178

16. Park SK, Mohr G, Yao J, Russell R, Lambowitz AM. Group II intron-like reverse transcriptases function in double-strand break repair. Cell 2022;185:1–18.

17. Nawrocki EP, Jones TA, Eddy SR. Group I introns are widespread in Archaea. Nucleic Acids Res. 2018;46:7970–7976. doi: 10.1093/nar/gky414.

18. Rogozin IB, Carmel, Csuros M, Koonin EV. Origin and evolution of spliceosomal introns. Biol. Dir. 2012;7:11. http://www.biology-direct.com/content/7/1/11.

19. Rogers SO, Integrated evolution of ribosomal RNAs, introns, and intron nurseries. Genetica 2019;147:103–119. doi.org/10.1007/s107009-018-0050-y.

20. Simmons MP, Bachy C, Sudek S, vanBare MJ, Sudek L, Ares M Jr, Worden AZ, Intron invasions trace algal speciation and reveal nearly identical Arctic and Antarctic *Micromonas* populations. Mol. Biol. Evol. 2015;32:2219–2235. doi:10.1093/molbev/msv122.

21. Bonocara RP, Shub DA. A likely pathway for formation of mobile group I introns. Curr. Biol. 2009;19;223–228. doi: 10.1016/j.cub.2009.01.033.

22. Tang TH, Rozhdestvensky TS, d’Orval BC, Bortolin M-L, Huber H, Charpentier B, Branlant C, Bachellerie J-P, Brosius J, Hüttenhofer A. RNAomics in Archaea reveals a further link between splicing of archaeal introns and rRNA processing. Nucleic Acids Res. 2002;30:921–930.

23. Hallsten U, Aspden JL, Rio DC, Rokhsar DS. A segmental genomic duplication generates a functional intron. Nature Comm. 2011;2:454. doi: 10.1038/ncomms1461.

24. Roca X, Sachidanandam R, Krainer AR. Intrinsic differences between authentic and cryptic 5’ splice sites. Nucleic Acids Res. 2003;31:6321–6333. doi: 10.1093/nar/gkg830.

25. Wright CJ, Smith CWJ, Jiggins CD. Alternative splicing as a source of phenotypic diversity. Nature Rev. Genet. 2022;23:697–710. doi: 10.1038/s41576-022-00514-4

26. Marasco LE, Kornblihtt AR. The physiology of alternative splicing. Nature Rev Molec Cell Biol. 2022. doi: 10.1038/s41580-022-00545-z.

27. Saudemont B, Popa A, Parmley JL, Necsulea A, Meyer E, Duret L. The fitness cost of mis-splicing is the main determinant of alternative splicing patterns Genome Biol. 2017;18;208. doi: 10.1186/s13059-017-1344-6.

28. Ward AJ, Cooper TA. The pathobiology of splicing, J. Pathol. 2010;220:152–163. doi:10.1002/path.2649.

29. Marchant DB, Chen G, Cai S, Chen F, Schafran P, et al. (45 additional authors), Dynamic genome evolution in a model fern. Nature Plants 2022;8:1038–1051. doi.org/10.1038/s41477-022-01226-7.

30. Rogers SO, Yan YH, LoBuglio KF, Shinohara M, Wang CJK. Messenger RNA intron in the nuclear 18S ribosomal DNA gene of deuteromycetes. Curr. Genet. 1993;23:338–342.

31. Shinohara ML, LoBuglio KF, Rogers SO. Group-I intron family in the nuclear ribosomal RNA small subunit genes of *Cenococcum geophilum*. Curr. Genet. 1996;29:377–387.

32. Harris L, Rogers SO. Splicing by an unusually small group I ribozyme. Curr. Genet. 2008;54:213–222.

33. Harris LB, Rogers SO. Evolution of small putative group I introns in the SSU rRNA gene locus of *Phialophora* species. BMC Res. Notes 2011;4:258–262.

34. Rogers SO. Evolution of the genetic code based on conservative changes of codons, amino acids, and aminoacyl tRNA synthetases. J. Theor. Biol. 2019;466:1–10.

35. Berget SM, Moore C, Sharp PA. Spliced segments at the 5ʹ terminus of adenovirus 2 late mRNA. Proc. Natl. Acad. Sci. U.S.A. 1977;74:3171–3175.

36. Chow LT, Gelinas RE, Broker TR, Roberts RJ. An amazing sequence arrangement at the 5ʹ ends of adenovirus 2 messenger RNA. Cell 1977;12:1–8.

37. Gilbert W. Why genes in pieces? Nature 1978;271:501.

38. Gao L, Altae-Tran H, Böhning F, Makarova KS, Segel M, Schmid-Burgk JL, Koob J, Wolf YI, Koonin EV, Zhang F. Diverse enzymatic activities mediate antiviral immunity in prokaryotes. Science 2020;369:1077–1084. doi: 10.1126/science.aba0372.32.

39. Kirchberger PC, Schmidt ML, Ochman H. The ingenuity of bacterial genomes. Annu. Rev. Microbiol. 2020;74:815–834. doi.org/10.1146/annurev-micro-020518-115822.

40. Koonin EV, Makarova KS. Mobile genetic elements and evolution of CRISPR-Cas systems: all the way there and back. Genome Biol. Evol. 2017;9:2812–2825. doi.org/10.1093/gbe/evx192.

41. Sakharkar MK, Chow VT, Kangueane P, Distributions of exons and introns in the human genome. In Silico Biol. 2004;4:387–393.

42. Xu X, Zhang J. Mammalian circular RNAs result largely from splicing errors. Cell Rep. 2021;36:109439. doi.org/10.1016/j.celrep.2021.109439.

43. Singh R. RNA–protein interactions that regulate pre-mRNA splicing. Gene Express. 2002;10:79–92.

44. Yoshihisa T. Handling tRNA introns, archaeal way and eukaryotic way. Frontier Genet. 2014. doi: 10.3389/fgene.2014.00213.

